# Simple spike dynamics of Purkinje cells in the macaque vestibulo-cerebellum reflect sensory prediction error

**DOI:** 10.1101/685461

**Authors:** Jean Laurens, Dora E. Angelaki

**Author notes:** Contact information: Dr. Dora E. Angelaki, Center for Neural Science, Meyer 901, New York University, NY 1003.

## Abstract

Theories of cerebellar functions posit that the cerebellum implements forward models for online correction of motor actions and sensory estimation. As an example of such computations, a forward model compensates for a sensory ambiguity where the peripheral otolith organs in the inner ear sense both head tilts and translations. Here we exploit the response dynamics of two functionally-coupled Purkinje cell types in the caudal vermis to understand their role in this computation. We find that one population encodes tilt velocity, whereas the other, translation-selective, population encodes linear acceleration. Using a dynamical model, we further show that these signals likely represent sensory prediction error for the on-line updating of tilt and translation estimates. These properties also reveal the need for temporal integration between the tilt-selective velocity and translation-selective acceleration population signals. We show that a simple model incorporating a biologically plausible short time constant can mediate the required temporal integration.

## Introduction

More than a century since the pioneering work of Ramon y Cajal (Cajal 1911), the cerebellum continues to represent a powerful model for understanding neural circuits. Its stereotyped anatomy (Palay & Chan-Palay 1976), its remarkably organized connectivity (Ruigrok 2011; Voogd 2011), and its profoundly tractable cellular identities (Eccles 1965; 1973) have motivated numerous recent advances in dissecting how cerebellar circuits are wired using modern molecular and optogenetic manipulations (Ankri et al 2015; Gao et al 2016; Guo et al 2014; 2016; Nguyen-Vu et al 2013; Witter et al 2016). In parallel to superb cellular and circuit organization discoveries, theory-driven studies have defined algorithmic computations likely performed by the cerebellar circuit. These computations extend beyond motor learning, into a modular organization for sensorimotor prediction and internal models (Wolpert et al 1998; Green & Angelaki 2010; Shadmehr et al 2010; Popa et al. 2012; 2013; 2016; 2017; Streng et al. 2018). However, little is currently known about how these computations map into the circuit. Thus, a major conceptual gap exists of how computational algorithms are mapped onto the canonical cerebellar circuit (Ito 2005).

One such internal model implemented by brainstem-cerebellar circuits merges signals from both vestibular end organs, the otoliths and semicircular canals, to resolve a sensory ambiguity (**Fig. 1A**) (Einstein, 1907): otolith afferents cannot distinguish linear acceleration (A) experienced during translations from gravitational acceleration (G) experienced during head tilt. Instead, otolith afferents encode the total gravito-inertial acceleration, GIA = G+A (**Fig. 1B**), thus responding identically to translational acceleration and tilt position (units: m/s^2^, or equivalently, ° of tilt). Theoretical (Mayne, 1974; Oman, 1982; Borah et al., 1988; Merfeld, 1995; Glasauer and Merfeld, 1997; Bos and Bles, 2002; Zupan and Merfeld, 2002; Laurens and Droulez, 2007; Laurens and Angelaki, 2011, 2017; Karmali and Merfeld, 2012; Lim et al., 2017) and experimental (Angelaki et al. 2004; Shaikh et al. 2005; Yakusheva et al. 2007; 2008; 2010; Laurens et al. 2013a,b; Dugué et al. 2017; Stay et al. 2019) studies have demonstrated that the brain resolves this ambiguity by using head rotation signals, originating from the vestibular semicircular canals, to track head movements relative to vertical, from which the gravitational component (G) can be estimated.

**Figure 1:**
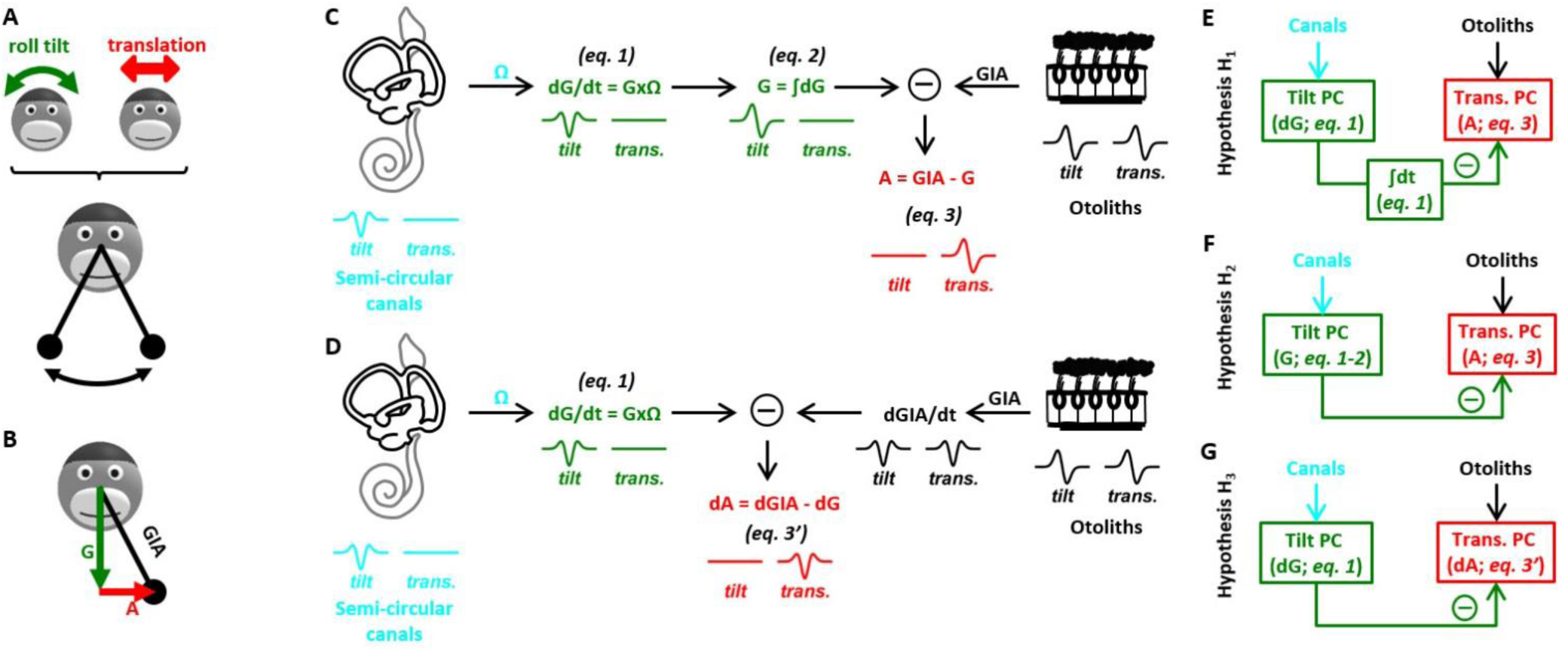
Internal model of head motion for resolving the tilt/translation ambiguity. (**A**) Illustration of the ambiguity: the otolith organs are analogous to a pendulum fixed to the head that swings identically during roll tilt and lateral translation. Thus, the otoliths detect both stimuli but do not discriminate them. (**B**) Illustration of the gravito-inertial force vector. (**C**) Simplified model of tilt/translation discrimination (from Laurens and Angelaki 2011). (**D**) Alternative architecture for tilt/translation discrimination. (**E-G**) Hypotheses (H_1_, H_2_, H_3_) of how internal model variables are represented in simple spike responses. The temporal waveforms shown are further detailed in **Fig. 1 Suppl. 1**, which shows a decomposition of the motion stimuli into dynamic components.

Although mathematical models of tilt-translation discrimination somewhat differ in their formulation (Mayne, 1974; Borah et al., 1988; Merfeld, 1995; Glasauer and Merfeld, 1997; Bos and Bles, 2002; Zupan and Merfeld, 2002; Laurens and Angelaki, 2011, 2017; Karmali and Merfeld, 2012), they all incorporate two salient computations (**Fig. 1C**): (1) the activity of semicircular canals, which encodes rotation velocity in an egocentric (head) reference frame (units: °/s) is spatially transformed (**Fig. 1C, *eq. 1***: the vectorial cross-product converts head-referenced rotation velocity signal (Ω) into a gravity-referenced tilt velocity signal, dG/dt); and (2) this canal-driven tilt signal must ‘combine’ with otolith afferent information, but the latter signals linear acceleration or tilt position relative to gravity. Often it has been assumed that the canal-driven, spatially-transformed signal must be temporally integrated (***eq. 2***: integration of tilt velocity signals into position) in order to estimate G, which is then subtracted from the otolith signal to compute linear acceleration (***eq. 3***). Note, however, that the brain might implement alternative but functionally equivalent computational schemes. In particular, ***eq. 3*** could be implemented in the velocity domain (**Fig. 1D, *eq. 3’***), implying a differentiation of the otolith-driven signal rather than an integration of the canal-driven signal.

Laurens et al (2013b) have indeed identified translation-selective and tilt-selective Purkinje cells as the neuronal correlates of the hypothesized tilt and translation signals. They have demonstrated that tilt-selective Purkinje cells encode spatially transformed signals (i.e. ***eq. 1*** or downstream) and that tilt- and translation-selective cells are functionally coupled (by ***eq. 3*** or ***eq. 3’***). However, the sinusoidal stimuli used in past experiments can’t resolve neuronal response dynamics. Therefore, whether tilt-selective neurons encode tilt (G) or tilt velocity (abbreviated here as ‘dG’), and whether translation-selective Purkinje cells encode linear acceleration (‘A’) or its derivative (abbreviated here as ‘dA’) is unknown. Distinguishing between these possibilities is a crucial step for understanding the computational algorithms implemented by central vestibular regions, and for identifying other components of the tilt/translation discrimination circuits.

In this study, we consider three alternative hypotheses, all of which would be consistent with the hypothesized computations: Tilt-selective cells may encode dG and translation-selective cells A (hypothesis H_1_, **Fig. 1E**). If this holds, then there is a functional need for temporal integration of the simple-spike signal of tilt-selective cells to implement ***eq. 2***, before it reaches translation-selective cells (***eq. 3***). This would suggest that another, yet unidentified, cell type, may encode a tilt signal (G). Alternatively, tilt-selective cells may encode G and translation-selective cells A (hypothesis H_2_, **Fig. 1F**). In this case, the integration (***eq. 2***) would occur upstream of tilt-selective Purkinje cells, or possibly in their dendritic tree. Finally, tilt-selective cells may encode dG and translation-selective cells dA (hypothesis H_3_, **Fig. 1G**), in which case the need for (***eq. 2***) would be eliminated.

Beyond understanding the tilt/translation disambiguation circuitry, discriminating between these hypotheses is also relevant for understanding how cerebellar networks implement sensorimotor internal models. There is growing evidence that not all types of error signals are carried by complex spikes (that lead to LTD of parallel fiber to Purkinje cell synapses; Marr, 1969; Albus, 1971; Ito and Kano, 1982; Ito, 2000). Additional error signals, which can cause plasticity in cerebellar and vestibular nuclei (Boyden et al. 2004; Ke et al. 2009), might be carried by the simple spike (SS) activity itself and encode feedback signals to optimize sensorimotor performance (Shadmehr et al. 2010; Popa et al. 2012; 2013; 2016; 2017; Streng et al. 2018). We recently implemented a Kalman filter model of self-motion sensation, where an internal model of head motion is continuously updated by feedback signals driven by sensory prediction errors (Laurens and Angelaki, 2017). This model, detailed further in Methods, predicts that feedback signals that update the internal estimates of tilt and translation should be proportional to tilt velocity and linear acceleration, respectively. Therefore, if Purkinje cells in the caudal vermis encode sensory prediction feedback signals, then the responses of tilt-selective Purkinje cells should correspond to ***eq. 1***, whereas the responses of translation-selective cells should correspond to ***eq. 3***. These Kalman filter model predictions favor hypotheses H_1_.

To distinguish among these three hypotheses (H_1_, H_2_, H_3_; **Fig. 1E-G**), we have recorded Purkinje cell simple spike (SS) activity using transient tilt, translation and tilt-translation stimuli that allow quantitative assessment of the response dynamics of tilt- and translation-selective Purkinje cells. A transient stimulus approach is necessary, as sinusoidal stimuli can’t resolve complex dynamic responses that do not follow linear systems properties (Angelaki and Dickman 2000; Dickman and Angelaki 2002; Laurens et al. 2017). The present results strongly support hypothesis H_1_, suggesting that Purkinje cell SS activity reflects sensory prediction errors. In particular, tilt-selective Purkinje cells may provide an on-line error signal for a forward model of how semicircular canal rotation signals can be mapped into an allocentric reference frame that governs spatial orientation and navigation in the terrestrial world (Laurens and Angelaki 2017).

## Results

### Experimental Findings

We recorded from NU Purkinje cells during transient tilt and translation stimuli with biphasic linear acceleration and Gaussian linear velocity profiles (σ = 250 ms), as illustrated in **Fig. 2A-E**. The tilt and translation stimuli were matched such that they activated the otoliths identically (**Fig. 2E**, first/second column; Angelaki et al. 2004; Shaikh et al. 2005; Yakusheva et al. 2007; 2008; 2010; Laurens et al. 2013a,b). During tilt-translation motion, tilt-driven and translation-driven otolith activation cancel each other (**Fig. 2E**, third column). Because the derivative of the biphasic tilt position (**Fig. 2B**, green) and linear acceleration (**Fig. 2C**, red) profiles follow a triphasic curve (e.g. tilt velocity in **Fig. 2D**, cyan), and because these signals ride on-top of a large spontaneous activity, the multiple temporal components of the models in **Fig. 1C** (i.e. G, A, GIA, dG/dt) as well as additional dynamic components (i.e. the integral of G and A, or the second derivative of G and A; **Fig. 1 Suppl. 1A**) can be distinguished.

**Figure 2:**
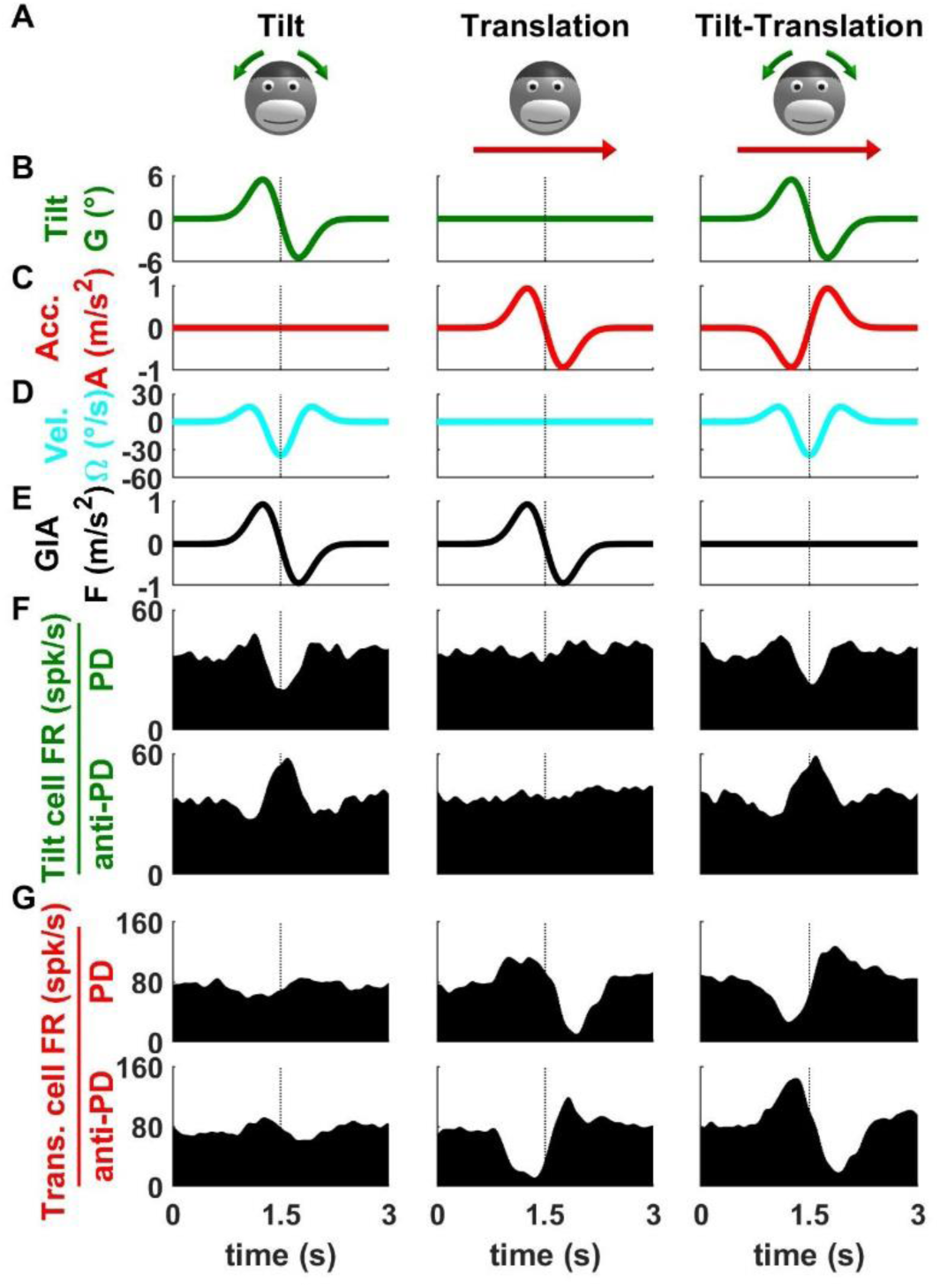
Example tilt- and translation-selective Purkinje cell responses during tilt, translation and tilt-translation. (**A**) Illustration of the motion stimuli. (**B-C**) Temporal profiles of the gravitational (i.e. tilt, B) and translational acceleration (C) component of the motion stimuli. (**D-E**) Temporal profiles of the physical variables sensed by the vestibular system: the tilt velocity (D) is detected by the semicircular canals and the gravito-inertial acceleration (GIA) (E) is detected by the otoliths. (**F-G**) Firing rate (FR) of a tilt-selective and translation-selective Purkinje cell. The upper and lower rows display the neuronal responses in the Preferred Direction (PD) and in the opposite direction (anti-PD), respectively. The PD is defined as the direction along which tilt-selective neurons increase their firing in response to positive tilt velocity and translation-selective neurons increase their firing in response to positive acceleration. Data shown in response to transient stimuli with σ = 250 ms.

Typical responses of tilt-selective and translation-selective Purkinje cells during the transient stimuli (with σ = 250 ms) are illustrated in **Fig. 2F**,**G**. During tilt, the example tilt cell exhibited a triphasic response modulation that was either proportional (preferred direction, PD; **Fig. 2F**, top) or inversely proportional (anti-PD; **Fig. 2F**, bottom) to tilt velocity (**Fig. 2D**, cyan). Here PD is defined as the direction along which firing rate is positively correlated with the stimulus; therefore the cell is inhibited during motion in its PD because tilt velocity is negative (**Fig. 2D**). The example tilt cell’s response resembles tilt velocity (the large peak/trough responses to tilt are flanked by smaller troughs/peaks) not only during tilt, but also during tilt-translation (**Fig. 2F**, left and right columns, respectively), but is negligible during translation (**Fig. 2F**, middle column). By contrast, the example translation cell modulates little during tilt (**Fig. 2G**, left), but responds vigorously to translation (**Fig. 2G**, middle) and tilt-translation (**Fig. 2G**, right). During translation along the cell’s PD (**Fig. 2G**, top), the cell exhibits a biphasic response whose dynamics follows the acceleration stimulus (**Fig. 2C**, red). The response reverses during motion along the anti-PD (**Fig. 2G**, bottom). Note that both tilt and translation Purkinje cells modulate during tilt-translation, when only the canals are dynamically modulated. This illustrates the fact that NU Purkinje cells receive convergent inputs from both sensors (Yakusheva et al. 2007; Laurens et al. 2013b).

These two example cells suggest that tilt Purkinje cells may follow tilt velocity (dG/dt), whereas translation Purkinje cells may follow linear acceleration (A), in support of hypothesis H_1_. We analyzed the transient responses of 30 NU Purkinje cells (3 macaques) which were specifically selected to be either tilt-selective (n=14) or translation-selective (n=16) following the criteria of Laurens and Angelaki (2013a,b). Note that cell classification was similar using transient and sinusoidal stimuli (**Table S1**).

We evaluated neuronal modulation by computing the difference in firing rate between motion in the PD and anti PD (**Fig. 3**). Note that this process cancels a quantitatively smaller omnidirectional component (**Fig. 3 Suppl. 1**) and only focuses on the direction-dependent responses. We measured each neuron’s peak-to-trough direction-dependent response during tilt and translation, as illustrated in the scatter plot of **Fig. 3A**). During translation, the responses of translation-selective cells were one order of magnitude larger than those of tilt-selective cells (389 spk/s/G, CI = [265-572] versus 39 spk/s/G, CI = [28-54]; p = 4.10^-6^, geometric mean and Wilcoxon sign rank test). In contrast, tilt- and translation-selective cells had comparable peak-to-trough modulation during tilt (tilt-selective cells: 151 spk/s/G, CI=[122-187]; translation-selective cells: 132 spk/s/G, CI=[94-184], p = 0.55). Thus, the range of response modulation amplitude during transient tilt and translation was remarkably similar to previous findings using sinusoidal stimuli (Laurens and Angelaki 2013b).

**Figure 3:**
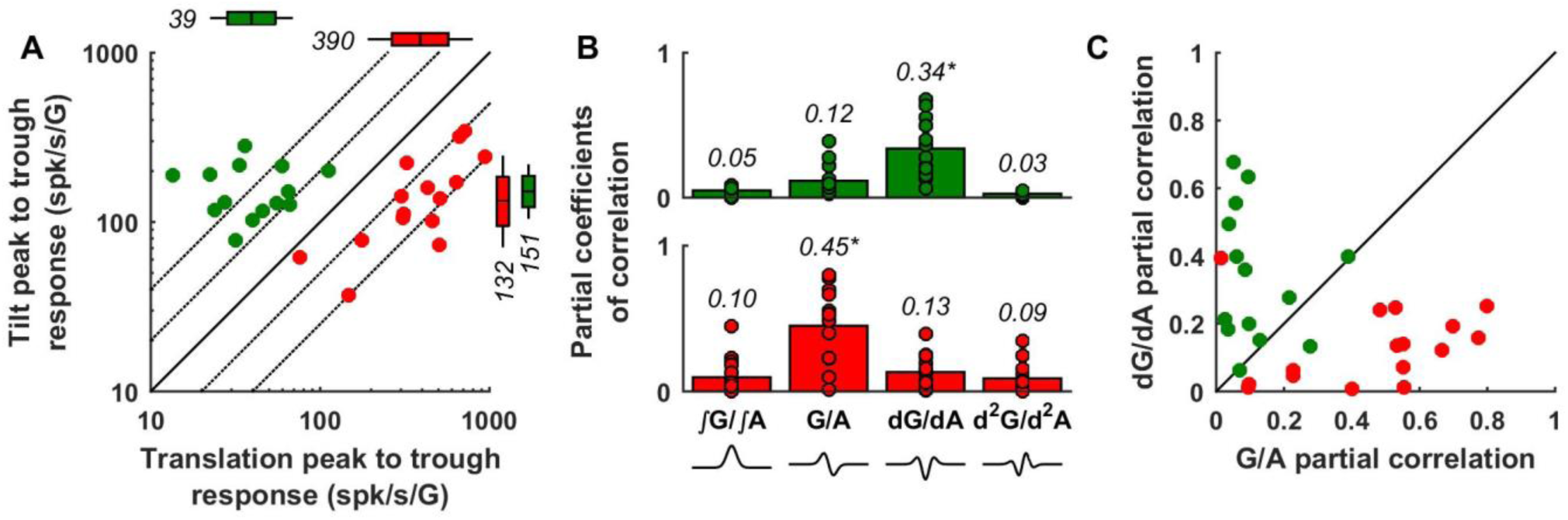
Dynamic response components of tilt (green) and translation (red) Purkinje cells. **(A)** Scatter plot of peak-to-trough response amplitude (in spk/s/G) during tilt and translation. The box and whisker plots indicate geometric mean (center of boxes), 95% confidence intervals (boxes) and standard deviation (whiskers). **(B)** Partial coefficients of correlation of the ∫G/∫A, G/A, dG/dA and d^2^G/d^2^A components (whose waveforms are illustrated at the bottom of the panel) in tilt (upper panel, green) and translation Purkinje cells (lower panel, red). Dots: individual cells, bars: population average. **(C)** Comparison of the partial correlation coefficients of biphasic (G/A) and triphasic (dG/dA) response components in tilt and translation selective Purkinje cells. Note that this analysis is agnostic to whether the cell encodes tilt or translation (because the G profile during tilt matches exactly the A profile during translation). Yet, it gives different answers for the two cell types – suggesting different dynamics. Data shown in response to transient stimuli with σ = 250 ms. All analyses are based on direction-dependent responses; summary of omnidirectional modulation responses is shown in **Fig. 3 Suppl. 1**.

Next we assessed which dynamic components are represented in neural responses. Our working hypotheses (H_1_, H_2_, H_3_) consider only two temporal components: (i) G (tilt position) and A (linear acceleration) – both of which have identical waveforms (**Fig. 1**; see also **Fig. 1 Suppl. 1**), and (ii) dG (abbreviation for dG/dt, tilt velocity) and dA (abbreviation for dA/dt, derivative of linear acceleration; i.e. jerk) signals – both of which also have identical waveforms (**Fig. 1**; see also **Fig. 1 Suppl. 1**). To characterize the cells’ response dynamics independently of their selectivity for tilt, translation or mixture thereof, we grouped motion variables with similar dynamics (i.e. G with A and dG with dA) and computed the partial coefficient of correlation of each pair (G/A and dG/dA) for each individual cell’s response. For generality, we included two additional dynamic components (∫G/∫A and d^2^G/d^2^A; **Fig. 1 Suppl. 1**) in the analysis. We found that the dG/dA component had the highest contribution to the responses of tilt cells (**Fig. 3B**, p<0.01, multiple paired Wilcoxon tests, Bonferroni correction), whereas the G/A component had the highest contribution in translation cells (**Fig. 3B**, p<0.01, multiple paired Wilcoxon tests, Bonferroni correction). In contrast, the partial correlation coefficients of the ∫G/∫A and d^2^G/d^2^A components were minimal. Thus, only the dG, dA, G and A components are considered in further analysis.

When plotted on a cell-by-cell basis, we found that the two cell types showed distinctly different response dynamics (**Fig. 3C**, green vs. red). Many tilt-selective cells clustered along the ordinate, and most (12/14, p = 0.002, paired Wilcoxon test) appear above the diagonal, indicating that the dG/dA profile dominates the responses of tilt-selective Purkinje cells. Considering that, by definition, tilt cells encode tilt, we conclude that tilt-selective cells carry predominantly a tilt velocity (dG) signal. Translation-selective cells clustered close to the abscissa and only one cell appeared above the diagonal (p = 0.0016), indicating that translation-selective cells carry acceleration (A) signals.

These conclusions are further illustrated in the average response profiles (**Fig. 4**; see also individual cell responses in **Fig. 4 Suppl. 1**). In line with the example cells in **Fig. 2F**, the average translation-selective cell exhibited a biphasic response profile that followed linear acceleration (**Fig. 4C**, red). The average tilt-selective cell exhibited a triphasic response profile that followed tilt velocity (**Fig. 4C**, green); although it displayed a slight asymmetry, where the second excitatory peak was attenuated compared to the first. This can be attributed to a small, but non-zero, G response, as shown in **Fig. 4D-F**. We plotted the G response component of tilt cells (**Fig. 4D**) as a function of their dG component. We found that both were correlated (p<10^-3^, bootstrap test), indicating that tilt-selective cells carry a G response component that is proportional to the dG component with about half the amplitude (slope = 0.47, CI = [0.27-0.79]). Plotting the average dG and G response components together (**Fig. 4E**) illustrates that the first peak of the G component (grey) tends to increase the first peak of the dG component (black), whereas the second peak of the G component reduces the last peak of the dG component. When these components are added (**Fig. 4F**, broken black line), this results in an asymmetrical profile that matches the average response profile of tilt cells (**Fig. 4F**, green, same as in **Fig. 4C**). This analysis, which reveals that tilt-selective cells encode primarily dG but also carry a weaker G component, is compatible with previous observations during sinusoidal motion at 0.5 Hz (Laurens et al. 2013b) where the response lagged tilt velocity by 36° (i.e. shifted towards tilt position). We repeated the same analysis for translation-selective cells (**Fig. 4G-I**). We found that these cells carry a small dA response (slope = -0.16, CI = [-0.24 to -0.06], p = 10^-3^), although this component was too small to alter the cell’s biphasic response profile markedly (**Fig. 4H**). In agreement, we observed (Laurens et al. 2013b) that the response phase of translation-selective cells was closely aligned with linear acceleration during sinusoidal motion.

**Figure 4:**
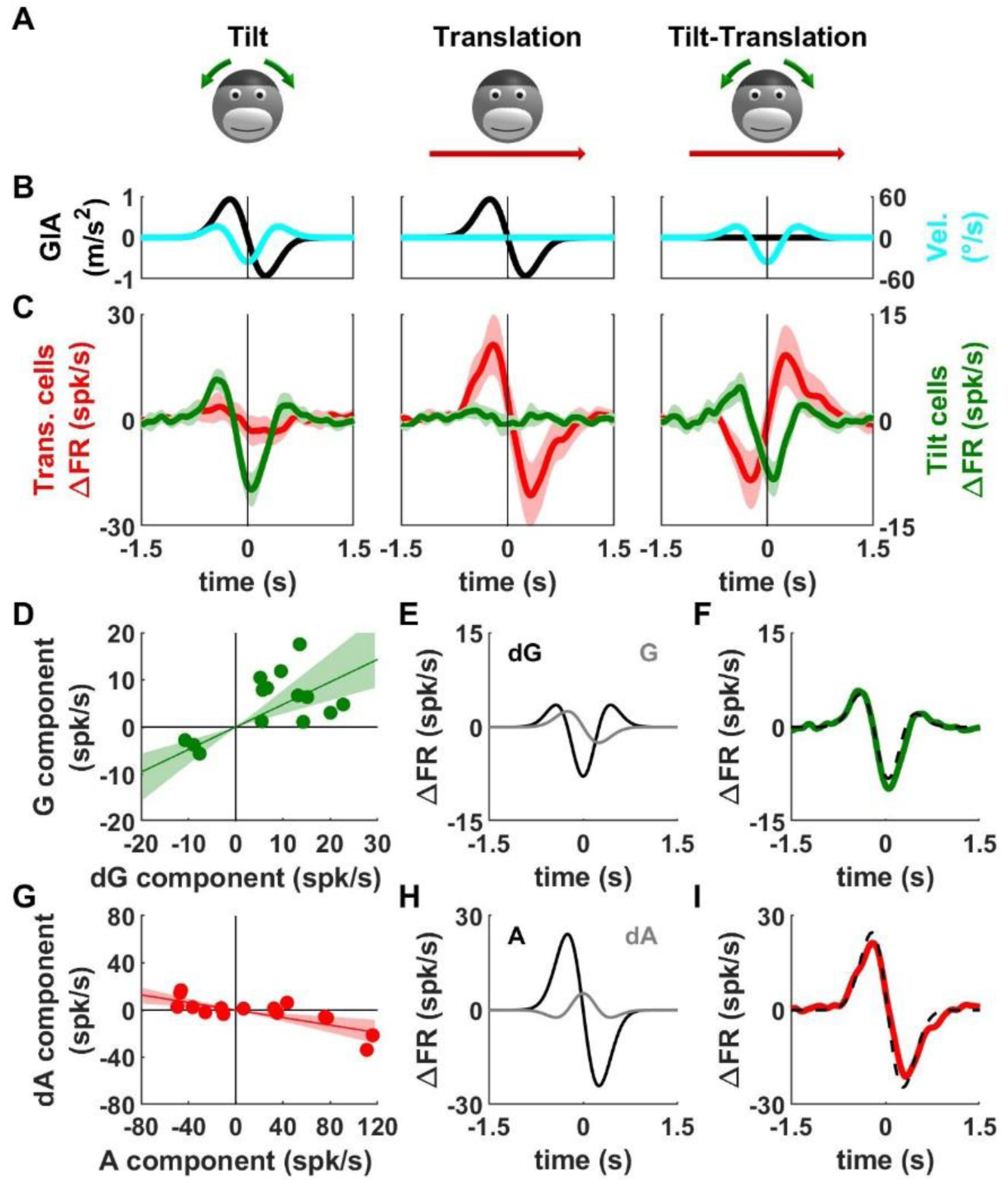
Average response profiles of tilt- and translation-selective cells. (**A-B**) Illustration of the stimuli (A) and sensory inputs (B), as in Fig. 2 (all individual cells are shown in **Fig. 4 Suppl. 1**). (**C**) Average direction-dependent response ΔFR (see Methods) of tilt-selective (green, right ordinate axis) and translation-selective (red, left ordinate axis) cells. The bands represent 95% CIs. Data shown in response to transient stimuli with σ = 250 ms. Summary of neuronal responses during transient motion of longer duration is shown in **Fig. 4 Suppl. 2. (D)** Correlation between the dG and G response components (defined as a “signed” peak to trough response amplitude, see Methods, *Composite model*) of tilt-selective cells. **(E)** Average dG (black) and G (grey) response components of tilt-selective cells (see Methods, *Composite model*). **(F)** Comparison between the sum of the dG and G components in (E) and the average response of tilt-selective cells in (C). **(G-H)** Analysis of the A and dA responses of translation-selective cells, as in (D-F).

Analyses of responses to a longer transient stimulus (σ = 500 ms) gave identical results (**Fig. 4 Suppl. 2**). In fact, other than a small but systematic increase in the gain of tilt cells (**Fig. 4 Suppl. 2E-F**), both sets of transient stimuli yield identical results.

### The dynamics of tilt- and translation-selective cells is consistent with feedback signals in an optimal model of head motion

There is now ample evidence that the brain separates gravity from linear acceleration (and processes self-motion information in general) by implementing a forward internal model of the vestibular organs (Borah et al. 1988; Merfeld 1995; Glasauer and Merfeld 1997; Merfeld et al. 1999; Angelaki et al. 1999, 2004; Laurens and Droulez 2007; Laurens and Angelaki 2011, 2017; Karmali and Merfeld 2012). This mechanism was recently formalized into a Kalman filter model (Laurens and Angelaki, 2017), where internal estimates of head motion (Ω, G and A) are used to predict vestibular afferent signals based on internal models of the semicircular canals and otolith organs. Differences between the predicted and actual afferent signals drive feedback loops that update the internal motion estimates.

In this schema, the internal model of the otoliths plays a central role in solving the gravito-inertial ambiguity, as outlined in **Fig. 5**. Rotation signals (derived from the internal model of the semicircular canals; see Laurens and Angelaki 2017 for details) are spatially transformed (***eq. 1***) and temporally integrated (***eq. 2***) to estimate G, as in **Fig. 1C**. The internal estimates of G and of linear acceleration (A) are fed into a forward internal model of the otoliths that predicts their activity (**Fig. 5**, ‘Otolith model’). Differences between the predicted and actual sensory inflow from the otoliths (GIA) results in feedback (**Fig. 5**, ‘Otolith feedback signals’) that corrects the internal estimates of acceleration (**Fig. 5**, ‘Acceleration’, red), tilt (**Fig. 5**, ‘Somatogravic tilt’, green, quantitatively minor here, see Methods) and rotation (‘Velocity’, cyan, see Laurens and Angelaki 2011, 2017 for details). During passive translations, the translation feedback closes a loop that implements ***eq. 3*** (see Laurens and Angelaki, 2017).

**Figure 5:**
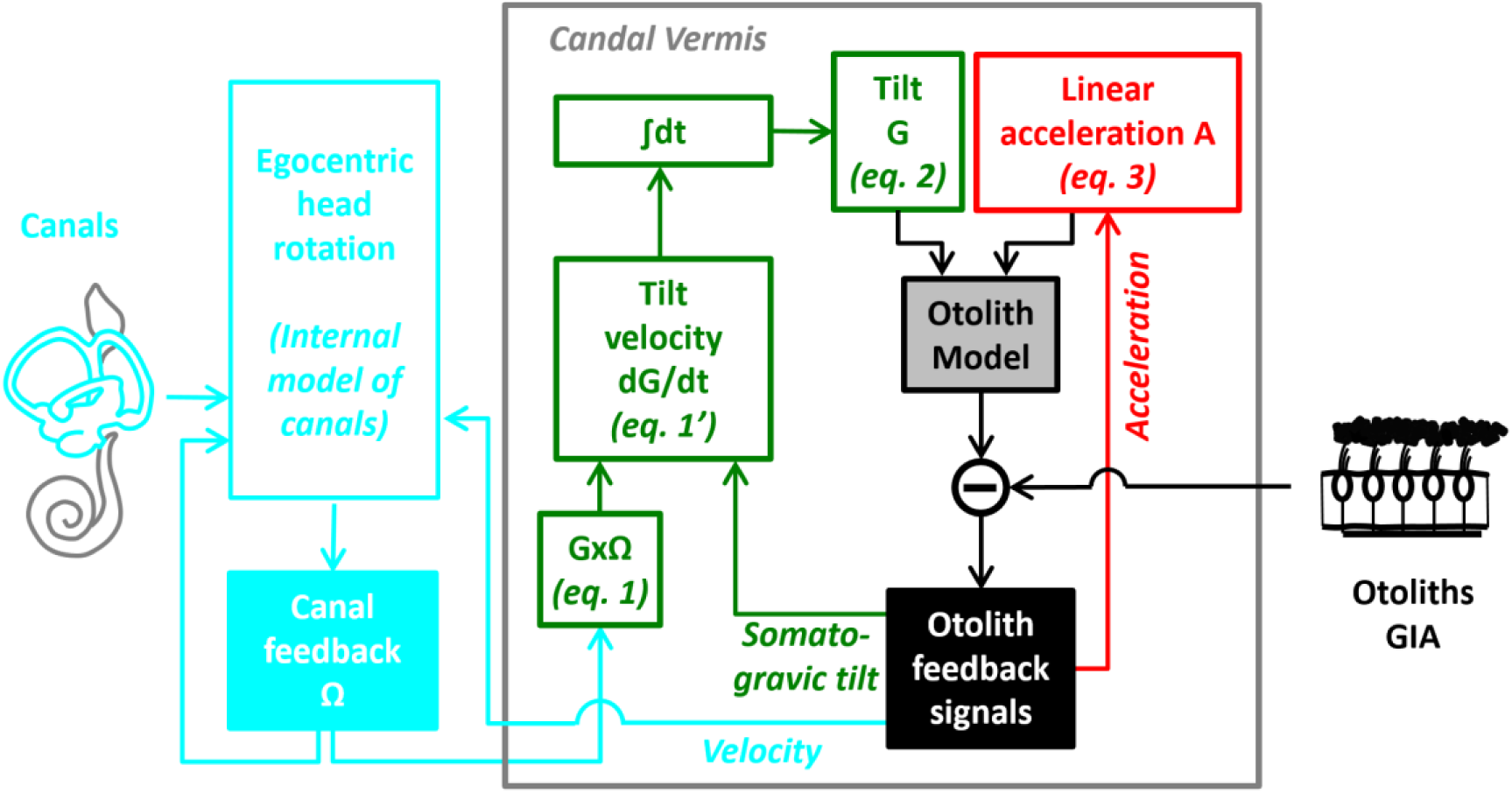
Kalman filter model of optimal processing of vestibular signals. Simplified internal model of vestibular information processing (from Laurens and Angelaki 2017). We propose that the caudal vermis compute internal estimates of tilt (green pathways) and translation (red pathways) that feed into an internal model of the otolith organs (black). During passive translations, the translation feedback closes a loop that implements ***eq. 3***, whereas the somatogravic feedback contributes a corrective component for ***eq. 1*** (see Methods and Laurens and Angelaki, 2017). Summary of the motion variables and equations are shown in **Fig. 5 Suppl. 1**.

In this study, we found that tilt-selective cells encode primarily tilt velocity. Furthermore, we found previously that they carry signals that correspond to the somatogravic feedback (Laurens and Angelaki, 2013b; see next section). We also found here that translation-selective cells encode linear acceleration. Thus, the responses of tilt- and translation-selective cells correspond to the properties of feedback pathways in an optimal model of vestibular information processing.

### A biologically plausible model of temporal integration

These experimental findings support hypothesis H_1_, where the output of tilt-selective Purkinje cells, which encode dG/dt, must get temporally integrated into a G signal (**Fig. 1C, *eq. 2***) before interacting with translation-selective Purkinje cells. This integration may be performed by a population of neurons (**Fig. 6A**, ‘Tilt Position’ neurons), functionally located between these Purkinje cell types. Yet, although (***eq. 2***) implies that this neuronal population should perform a perfect integration, this operation may not be biologically plausible. Instead, the hypothesized neuronal population may perform a leaky integration, with a time constant of ∼1s, and therefore integrate canal-driven rotation signal accurately at high frequencies only. At low frequencies, this population’s activity may be sustained by otolith-driven somatogravic feedback signals conveyed by tilt-selective cells. To test this scheme quantitatively, we simulated a network model (see Methods) during transient tilt and translation as well as static tilt (**Fig. 6B**,**C**). As shown in **Fig. 6D-F**, simulations agree with experimental data. First, tilt-selective Purkinje cells (**Fig. 6D**, green) follow tilt velocity (**Fig. 6C**, cyan) during tilt, and their simulated response is reduced during static tilt (since tilt velocity is null) and translation (where only a faint response, driven by the somatogravic feedback, is observed). In contrast, the intermediate neuronal population (**Fig. 6E**, green) responds in phase with tilt position (**Fig. 6B**, green), including static tilt (although with a smaller gain; 0.72 compared to dynamic tilt), and have reduced responses during translation. Finally, translation-selective Purkinje cells (**Fig. 6F**, red) follow linear acceleration (**Fig. 6B**, red) and have reduced responses during all tilt protocols. Thus, our simulations confirm that G signals may be computed by a leaky integration with a time constant of 1s, in conjunction with the somatogravic effect, which provides a steady-state tilt signal.

**Figure 6:**
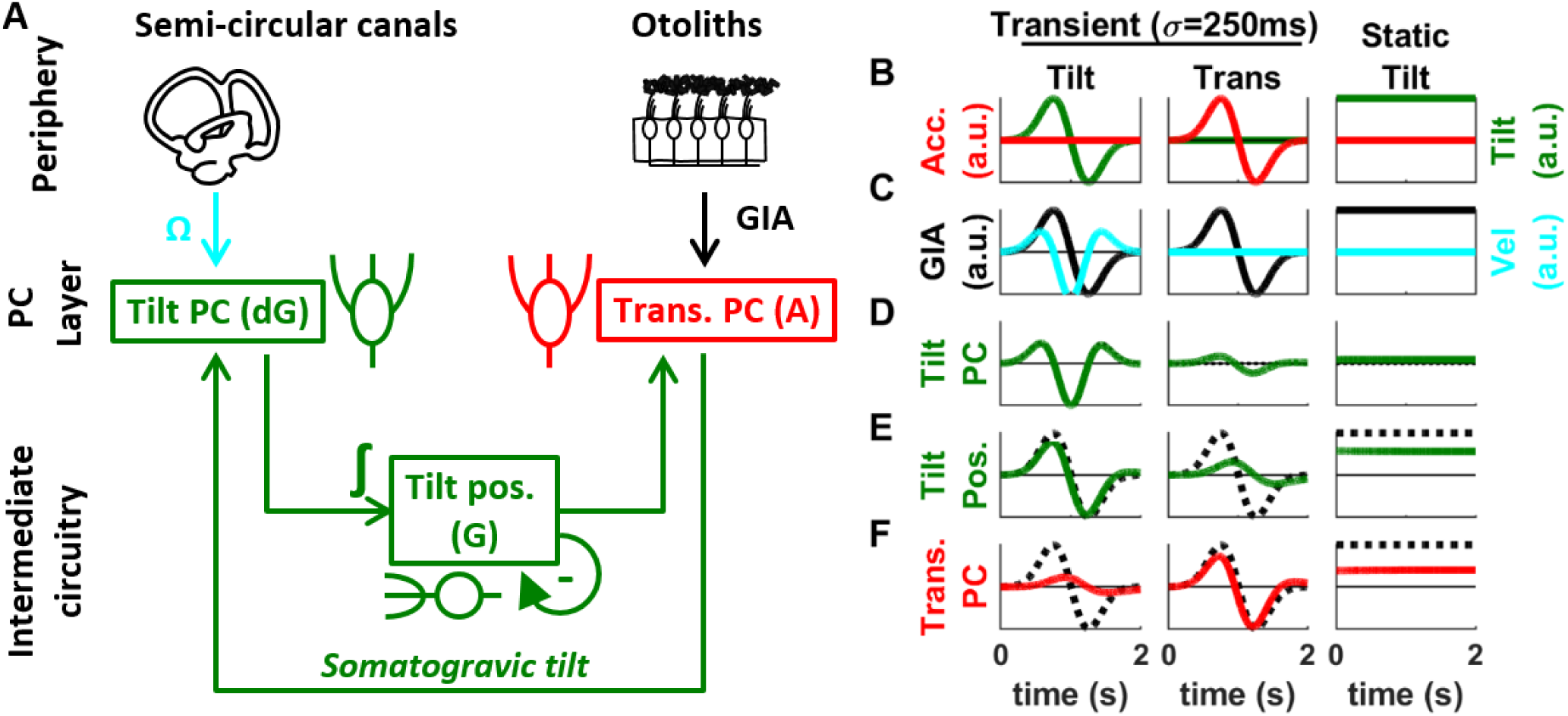
Modeling hypothesis H_1_ into a simplified neural circuit. **(A)** Overview of the proposed neuronal circuitry. **(B-F)** Simulations of the network during tilt and translation. **(B)** Motion variables (tilt and linear acceleration). **(C)** Sensory variables (GIA and tilt velocity). **(D)** Simulated response of tilt PCs (green). Note that the simulated cells exhibit a faint response during translation **(E)** Simulated response of a neuron encoding tilt position (green). The broken black line represents the GIA. Note **(F)** Simulated responses of a translation-selective PC (red). The broken black line represents the GIA. All variables are expressed in arbitrary units.

## Discussion

We have shown that tilt- and translation-selective Purkinje cells differ in response dynamics: tilt-selective cells encode primarily tilt velocity, whereas translation-selective cells encode linear acceleration. These dynamics are consistent with the notion that the simple spike response of Purkinje cells encode feedback signals that derive from sensory predictions errors and are computed by a forward internal model of the vestibular organs (**Fig. 5**; Laurens and Angelaki 2017).

Laurens et al. (2013b) has shown that the sum of the population responses of tilt and translation-selective Purkinje cells sum to the GIA. Furthermore, local gabazine injection into the cortex converts translation-selective Purkinje cells into GIA-coding cells (Yakusheva et al., 2013). Thus, it has been proposed that translation-selective Purkinje cells generate their responses through convergence of otolith (GIA) and the tilt-selective Purkinje cell population signals (Laurens et al. 2013b). Combined with the present findings, we conclude that the output of tilt Purkinje cells should be temporally integrated before relayed to translation-selective Purkinje cells. Simulations (**Fig. 6**) suggest that this may be accomplished by a simple neuronal circuit performing a leaky integration with a biologically plausible time constant.

The framework of internal models has been the dominant theory of vestibular processing in the past decades (Mayne, 1974; Oman, 1982; Borah et al., 1988; Merfeld, 1995; Glasauer and Merfeld, 1997; Bos and Bles, 2002; Zupan and Merfeld, 2002; Laurens, 2006; Laurens and Droulez, 2007, 2008; Laurens and Angelaki, 2011, 2017; Karmali and Merfeld, 2012; Lim et al., 2017) and is closely related to the framework used to model motor control and adaptation (Wolpert et al., 1995; Körding and Wolpert, 2004; Todorov, 2004; Chen-Harris et al., 2008; Berniker et al., 2010; Berniker and Körding, 2011; Franklin and Wolpert, 2011; Sağlam et al., 2011, 2014). Initially supported by behavioral studies using passive motion stimuli (e.g. Merfeld et al. 1993, 1999; Laurens et al. 2010, 2011), the implementation of internal models has been confirmed by neurophysiological experiments of tilt/translation discrimination (Angelaki et al. 2004; Shaikh et al. 2005; Yakusheva et al. 2007; 2008; 2010; Laurens et al. 2013a,b; Dugué et al. 2017; Stay et al. 2019) and active head movements (Roy and Cullen, 2004; Cullen et al., 2011; Cullen, 2012; Carriot et al., 2013; Brooks and Cullen, 2013,2014; Brooks et al., 2015). Laurens and Angelaki (2017) have formulated a Kalman filter to model neuronal responses in the vestibular nuclei, cerebellar cortex and deep cerebellar nuclei during both active and passive motion (shown in simplified form in **Fig. 5A**). The present study demonstrates that the SS response dynamics of tilt- and translation-selective Purkinje cells reflects tilt velocity and translation feedback signals predicted by the Kalman filter. This finding supports the hypothesis that the SS activity of Purkinje cells carry sensory prediction error signals, a critical component of a dynamical control framework supporting optimal sensorimotor functions (Shadmehr et al. 2010; Popa et al. 2012; 2013; 2016; 2017; Streng et al. 2018).

The Kalman filter also predicts that feedback signals, and consequently the activity of tilt- and translation-selective cells, should be profoundly attenuated during active tilt and translation, similar to neuronal responses measured in the vestibular nuclei, fastigial nuclei and cerebellar cortex (Roy and Cullen, 2004; Cullen et al., 2011; Cullen, 2012; Carriot et al., 2013; Brooks and Cullen, 2013,2014; Brooks et al., 2011; Lee et al. 2015; Dugué et al., 2017). In agreement with this prediction, one study (Lee et al. 2015) conducted when rats learn to balance on a swing indicates that Purkinje cells in various lobules (V to X) of the cerebellar vermis encode tilt velocity during external perturbations, but not learned active movement. Interestingly, the Kalman filter predicts that the central estimate of tilt, which may be carried by cortical interneurons (see next paragraph), should not be attenuated during active tilt. Future recording studies of cerebellar interneurons should investigate these predictions further.

Although tilt-selective Purkinje cells encode predominantly tilt velocity, we found that they carry a smaller but consistent tilt position component (**Fig. 4D-F**). This finding is consistent with Laurens et al. (2013b), which reported response phase shifted by 36° towards tilt position during sinusoidal tilt at 0.5Hz. Thus, tilt-selective Purkinje cells may themselves be within the dynamic system that generates the tilt position signal, although their responses remain closer to tilt velocity than position.

Our results suggest that the simple-spike output of tilt-selective cells may be temporally integrated by an intermediate neuronal type. One possibility is that this temporal integration occurs outside the cerebellar cortex, such that G signals reach translation-selective Purkinje cells through mossy fiber projections from the vestibular nuclei. We also consider a more parsimonious explanation based on the recently discovered Purkinje axon collaterals onto the cerebellar cortex (Guo et al. 2016; Witter et al. 2016), such that the temporal integration may involve granular layer interneurons, e.g., unipolar brush cells (UBCs) and/or granule/Golgi cells. That UBCs may be involved is supported by both in-vitro (van Dorp and De Zeeuw, 2014; 2015; Locatelli et al. 2013) and in-vivo (Kennedy et al. 2014) findings. UBCs receive extensive synaptic contacts from a single mossy fiber rosette from either vestibular afferents or vestibular nuclei (Barmack et al. 1992; Diño et al. 2001; Jaarsma et al. 1996) and exert a powerful excitatory action onto multiple granule cells and other UBCs (Dino et al. 2000; Nunzi and Mugnaini, 2000). This highly specialized configuration is thought to facilitate prolonged entrapment of glutamate and broaden the temporal window of activation, thus facilitating temporal transformations (Zampini et al. 2016).

Of particular interest may be calretinin-positive UBCs, which are specifically found in the NU (Kim et al. 2012; Sekerkova et al. 2014) and receive vestibular afferent mossy fibers (Dino et al. 2000). Alternatively, it could be that the G signal is found in other UBC types or granule interneurons, which perform multimodal integration (Arenz et al. 2008; 2009; Chabrol et al. 2015; Ishikawa et al. 2015) and receive unusually massive collaterals from Purkinje cells in the NU (Guo et al. 2016). Feedback connections from the cerebellar nuclei to the cerebellar cortex may also contribute. For example, some cerebellar nuclei neurons send collaterals back to the cortex contacting granule and Golgi cells (Ankri et al. 2015; Gao et al. 2016; Houck and Person, 2015). Furthermore, glutamatergic neurons in the nuclei, in addition to projecting to various premotor and associative regions of the brain, send axonal collaterals to form mossy fiber-like terminals contacting granule and Golgi cell dendrites (Houck and Person, 2015). More recently, an inhibitory nucleo-cortical feedback loop was established. Ankri et al. (2015) found that GABA-glycinergic nuclei neurons form an extensive and divergent plexus of axons, which contact Golgi cells in the cerebellar granular and molecular layers. Notably, neither rosette-like terminals nor evidence of contacts within cerebellar glomeruli was found. This indicates that they differ both in shape and location from the excitatory mossy fibers and the glutamatergic nucleo-cortical fibers, both of which form rosette-like terminals within the glomeruli (Tolbert et al. 1978; Hámori et al. 1980; Batini et al. 1992; Houck and Person, 2015). It is important that future studies test these hypotheses explicitly.

## Acknowledgements

The work was supported by NIH grant DC004260. The authors would like to acknowledge Hui Meng’s contributions to the recordings.

## Methods

### Animals

Three male rhesus Macaques, aged 3, 4 and 9 years, were used in the study. Animals were pair-housed in a vivarium under normal day/night cycle illumination. Animals were implanted under isoflurane anesthesia with a circular delrin ring to immobilize the head, scleral search coils to measure eye movements and a delrin platform for neural recordings. Experimental procedures were in accordance with US National Institutes of Health guidelines and approved by the Animal Studies Committee at Washington University in St Louis (approval n°20100230) and Baylor College of Medicine (protocol n°AN-5795).

### Experimental setup and neuronal recordings

Experimental procedures were similar as in previous studies (Yakusheva et al. 2007; Laurens et al. 2013a,b). Primates sat comfortably in a primate chair that was installed in the center of a 3-axis rotator mounted on a linear sled (Acutronics Inc, Pittsburg, PA) such that the three rotation axes intersected at the center of the head. Neurons were recorded extracellularly using epoxy-coated tungsten microelectrodes (5 or 20 MΩ impedance; FHC, Bowdoinham, ME), acquired at 33kHz using a data acquisition board (1401, Cambridge Electronic Design, Cambridge, UK) and stored for offline analysis. Spike sorting used a custom Matlab script (MathWorks) by manually clustering spikes based on spike amplitude and principal components analysis.

The location of lobules X and IX of the caudal vermis was determined based on stereotaxic coordinates as well as the location of the abducens nucleus. Recordings were performed in the Purkinje cell layer where complex spikes activity could be observed online. Complex spikes were further identified offline in 33/46 cells, and simple spike activity was observed to pause for at least 15ms in 30/33 cells.

### Experimental protocol

Once neural activity was isolated, we used sinusoidal tilt and translation stimuli (0.5Hz, 0.2G amplitude; as in Shaikh et al. 2005; Yakusheva et al. 2007; Laurens et al. 2013a,b) to determine online whether the cell responded preferentially to tilt or translation using motion along multiple axes (naso-occipital, inter-aural or intermediate). Because our focus was on tilt- or translation-selective neurons, only cells with a clear modulation during tilt-translation were further tested using a series of transient stimuli along the cell’s preferred direction.

The transient motion profiles were generated by computing the derivative of a Gaussian function with standard deviation σ = 250ms. This resulted in a biphasic signal that was scaled to an amplitude of ±5.6° to generate the tilt position stimulus, and to an amplitude of ±0.93m/s^2^ to generate the linear acceleration stimulus. A tilt-translation stimulus was created by applying tilt and translation stimuli simultaneously so that the resultant gravito-inertial acceleration was null. Each stimulus type (tilt, translation and tilt-translation) was applied 15 times in two opposite directions. Longer duration stimuli were generated by setting σ to 500ms, and the peak tilt and linear acceleration amplitudes to ±9.8° and ±1.67m/s^2^.

### Sample size

In line with standard practices in extracellular studies in non-human primates, we aimed at collecting a sample of over 40 neurons in over 2 animals. Our final sample includes 46 neurons in 3 animals.

### Data analysis

Neuronal responses were analyzed using a linear model schematized in **Fig. 1 Suppl. 1**. We computed a peri-stimulus time histogram (time t ranging from -2s to 2s by increments of 12 ms, total of 333 bins) for each stimulus type (Tilt, Translation, Tilt-Translation) and motion direction (positive or negative).

As shown by Laurens et al. (2017), neural responses to translation are dynamically complex, consisting of both spatially-tuned (direction-selective) and spatially-untuned (omnidirectional) components. For illustration purpose (and separately from the linear regression analysis described below), the direction-selective components can be visualized by computing the difference between the firing rates measured during motion in both direction (ΔFR = (FR_PD_ - FR_Anti-PD_)/2), where FR_PD_ and FR_Anti-PD_ are the cell’s firing along its preferred motion direction (PD) or in the opposite direction (Anti-PD). A cell’s PD refers to its direction-selective modulation, and is defined as the direction along which tilt-selective neurons increase their firing in response to positive tilt velocity and translation-selective neurons increase their firing in response to positive acceleration. The omnidirectional component can be visualized by computing the average firing across both directions.

To characterize response dynamics, we used multiple linear regression. Specifically, we decomposed tilt into 4 dynamic components (**Fig. 1 Suppl. 1**): tilt position (G), tilt velocity (dG/dt, abbreviated dG), tilt acceleration (d^2^G/dt^2^, abbreviated d^2^G) and the integral of tilt position (∫G.dt, abbreviated ∫G). Likewise, we decomposed linear acceleration into (∫A, A, dA, d^2^A, i.e. linear velocity, acceleration, jerk and jerk derivative respectively; **Fig. 1 Suppl. 1**). As shown by Laurens et al. (2017), neural response may include omnidirectional components, where cells respond identically (e.g. by an increase in firing rate) irrespective of motion direction. To quantify these response components, we added 8 additional regressors (∫G^O^, G^O^, etc…) which were identical to their counterpart (∫G, G, etc…, also referred to as “direction-dependent”) but did not reverse sign for opposite motion directions (**Fig. 1 Suppl. 1**, “Omnidirectional motion variables”). Next, we performed a series of linear regressions where all peri-stimulus time histograms (along all directions, i.e. we didn’t extract the direction-selective and omnidirectional components prior to this analysis) were simultaneously fitted with either all or a subset of theses 16 variables.

#### Composite model

The first regression, which included all variables, the *composite* model, followed the equation:

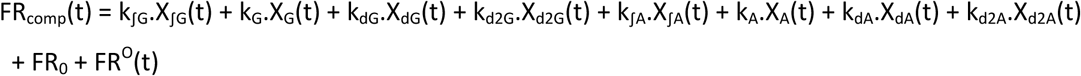

in this equation, FR_0_ is the cell’s baseline firing rate, and the omnidirectional motion variables have been grouped in a variable FR^O^(t):

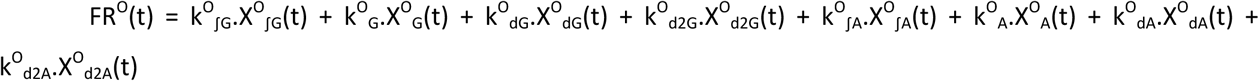

The regression coefficients (k_∫G_, k_G_, k_dG_, etc…) were used to evaluate the neurons’ response gain to G, dG, etc. Note that the composite model included 16 temporal variables that are all linearly independent (therefore the system was not overdetermined) and are all statistically orthogonal when only tilt and translation motion are considered. This property ensures that the composite model is not prone to overfitting. Note also that the purpose of this analysis was not to demonstrate that neuronal responses could be fitted accurately (which would not be very remarkable, considering the large number of variables used in the model), but to investigate which variables contributed to the neuron’s response.

The neuronal response gains may not be directly compared across dynamic components since they are expressed in different units (e.g. spk/s/G for A and G, spk/s/(G/s) for dA and dG). To convert them to identical units, we scaled the regression coefficients (k_∫G_, k_G_, k_dG_, etc…) by the peak to trough amplitude of the motion variables (X_∫G_, X_G_, X_dG_…), resulting in “signed” peak-to-trough response amplitudes (in spk/s) that can be compared across dynamic components (**Fig. 4D, G**). In **Fig. 4E, H**, the temporal profiles of tilt-selective (or translation-selective) cells are computed as the average values of |k_dG_| and |k_G_| (|k_dA_| and |k_A_| respectively) multiplied by the temporal profiles of X_dG_ and X_G_ (X_dA_ and X_A_ respectively).

#### Partial correlation analysis

In order to evaluate how well a single motion variable or a group of variables (e.g. ∫G and ∫A) contributes to a neuron’s response, we re-fitted the firing rate after eliminating the motion variable (or group of variables). The partial coefficient of determination (pR^2^) of this group of variables is computed as:

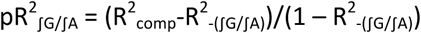

Where R^2^_-(∫G/∫A)_ is the coefficient of determination after k_∫G_.X_∫G_(t) and k_∫A_.X_∫A_(t) are removed from the composite model.

Note that this computation was also done for pair of variables with identical dynamic components (∫G and ∫A; G and A; etc) in order to quantify the cells’ dynamics irrespectively of whether cells preferentially responded to tilt, translation or GIA (**Fig. 3B**,**C**). In addition, we also computed the partial R^2^ of all direction-dependent components together (pR^2^_direction-dependant_), as well as the partial R^2^ of all omnidirectional components (pR^2^_omnidirectional_) (**Fig. 3 Suppl. 1F**).

#### Neuronal response classification

Following a similar approach as in Laurens and Angelaki 2013b, we performed additional regressions based on subset of motion variables to classify the cells as tilt-selective, translation-selective, GIA-selective or composite. We fitted FR(t) with simpler models that assume that the neuron responds exclusively to tilt, translation or the GIA:

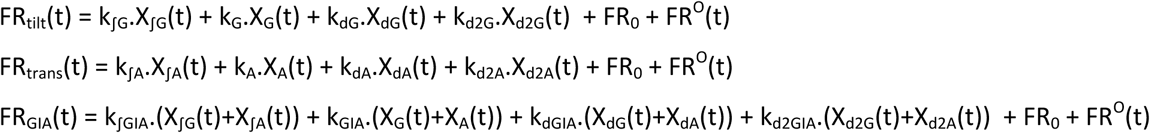

The quality of each model’s fit was evaluated by computing coefficients of determination R^2^_tilt_, R^2^_trans_, R^2^_GIA_. A neuron was classified as tilt-, translation- or GIA selective if its R^2^ was significantly (p<0.01) higher than the R^2^ of the two other models; the p-value was computed using a bootstrap procedure (as in Laurens and Angelaki 2013b). If no component was significantly higher than the others, the neuron was classified as composite. Any neuron where pR^2^_direction-dependant_ <0.3 was classified as non-responsive.

Note that cells were classified based on their “direction-dependent” response alone. Indeed, although the omnidirectional response component may also encode tilt, translation or GIA, these response components were captured by the term FR^O^(t) which was included in all the models above. Therefore, these models did fit the omnidirectional response equally well, and differed only by their ability to fit the direction-dependent responses.

A similar procedure was performed to classify omnidirectional responses (**Fig. 3S1, Suppl. Table 2**).

#### Kalman filter model

The Kalman filter model in Laurens and Angelaki 2017 computes optimal estimates of rotation velocity, tilt and translation during active and passive motion. In **Fig. 5**, we have outlined a simplified version, which is restricted to estimating tilt and translation during passive movement. This model implements the computations outlined in **Fig. 1C**, i.e. ***eq. 1-3*** (see Laurens and Angelaki 2017 for details). Note that the model incorporates an otolith feedback that updates the tilt estimate (somatogravic feedback), see **Fig. 5 Suppl. 1**. In the absence of canal inputs, i.e. Ω=0, ***eq. 1’*** implements a low-pass filter. This corresponds to a well-known illusion where tilt sensation follows otolith signal at low frequencies (Graybiel, 1952). This term has only a minor contribution for the stimuli used, as shown in **Fig. 5 Suppl. 1** (compare the simulations of ***eq. 1*** and ***eq. 1’***).

#### Neuronal network simulations

We simulated a putative cerebellar circuitry where the firing rate of tilt-selective Purkinje cells (FR_TiltPC_), interneurons (FR_TiltPos_) and translation-selective Purkinje cells (FR_TransPC_) implement dG/dt, G and A, respectively. We assumed that tilt-selective Purkinje cells compute an optimal estimate of tilt velocity by implementing ***eq. 1’*** (**Fig. 5**). Accordingly, we model these cells using eq. 1” below, which is a direct transcription of ***eq. 1’***:

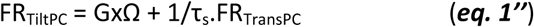

Ideally, neurons that encode tilt should implement ***eq. 2*** by integrating tilt velocity signals provided by tilt-selective Purkinje cells inputs to compute gravity. However, ***eq. 2*** (G=∫dG.dt) stipulates that dG signals should be integrated perfectly, but this might not be practically feasible. In fact, because the integration of semicircular canal signals into an internal estimate of G occurs at high frequencies (as shown in Laurens et al. 2013b), whereas the somatogravic feedback dominates this estimate at low frequencies, we reasoned that a leaky integrator would approximate ***eq. 2*** closely:

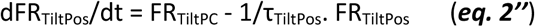

Finally, we assume that translation-selective cells implement ***eq. 3***, and accordingly we model these cells using eq. 3” below, which is a direct transcription of ***eq. 3***:

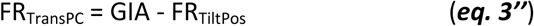

We set τ_s_ = 0.5s and τ_TiltPos_ = 1s and simulated the network’s dynamics during transient motion stimuli as well as static tilt (**Fig. 6B-F**).

---

**Figure 1 Supplement 1:**
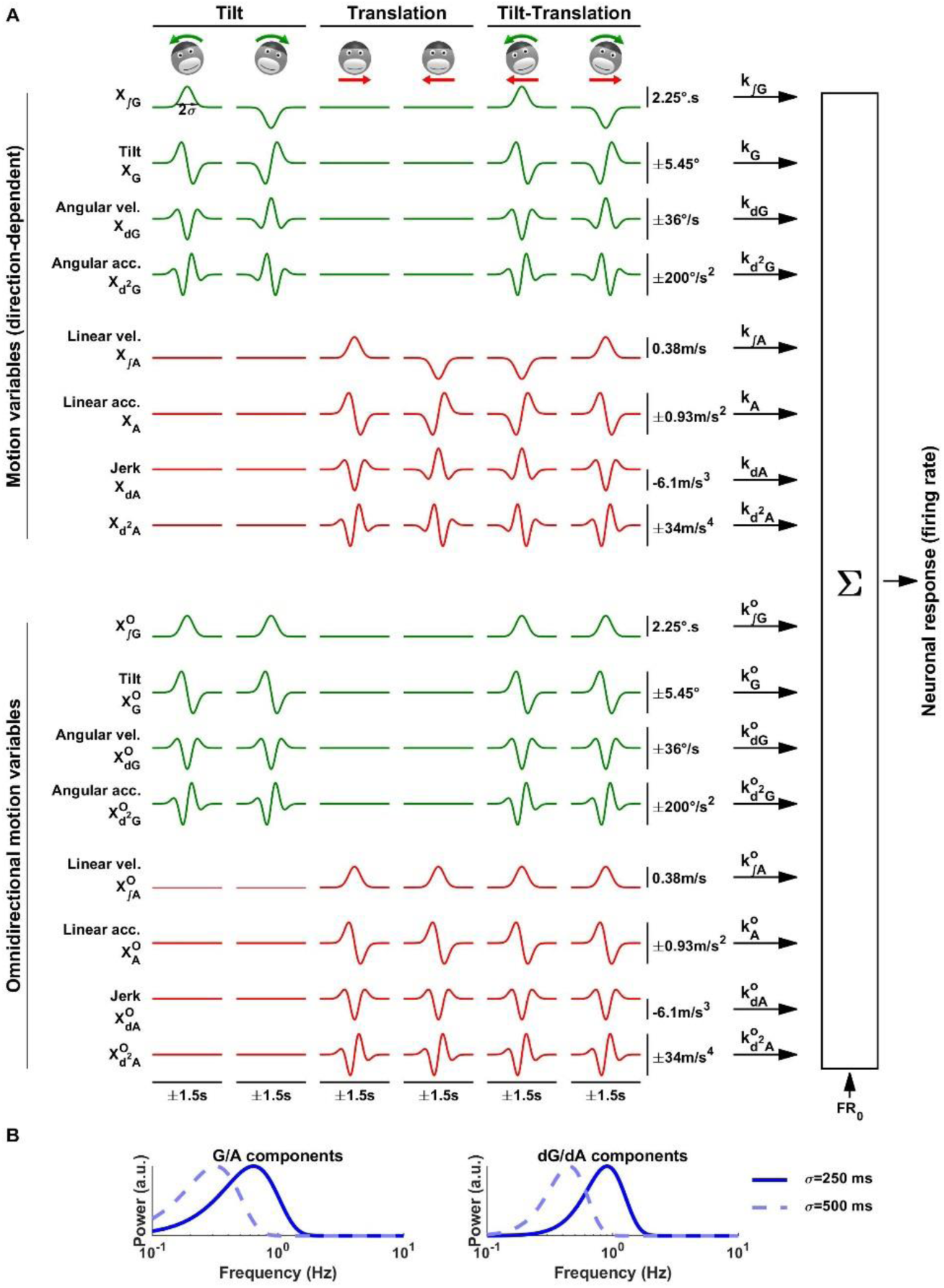
Dynamics of the transient motion stimuli. **(A)** Decomposition of the motion stimuli and neuronal responses into dynamic components. Motion stimuli are illustrated on top (assuming lateral motion; similar stimuli were also applied in the forward/backward or intermediate directions). Arrows represent the direction of the first phase of the biphasic tilt or linear acceleration profiles. Upper half: Motion variables. Tilt (G) and linear acceleration (A) follow biphasic profiles that can be integrated into ∫G and ∫A or derivated into dG/dA and d^2^G/ d^2^A. The corresponding temporal profiles during tilt, translation and tilt-translation in both directions (with σ=250ms) are represented. Lower half: Omnidirectional motion variables. We define omnidirectional variables that follow the same dynamics as the motion variables, but whose sign doesn’t reverse when the direction of motion is changed. Neuronal responses are modeled as linear combinations of these components. **(B)** Power spectra of the G/A and dG/dA components.

**Figure 3 Supplement 1:**
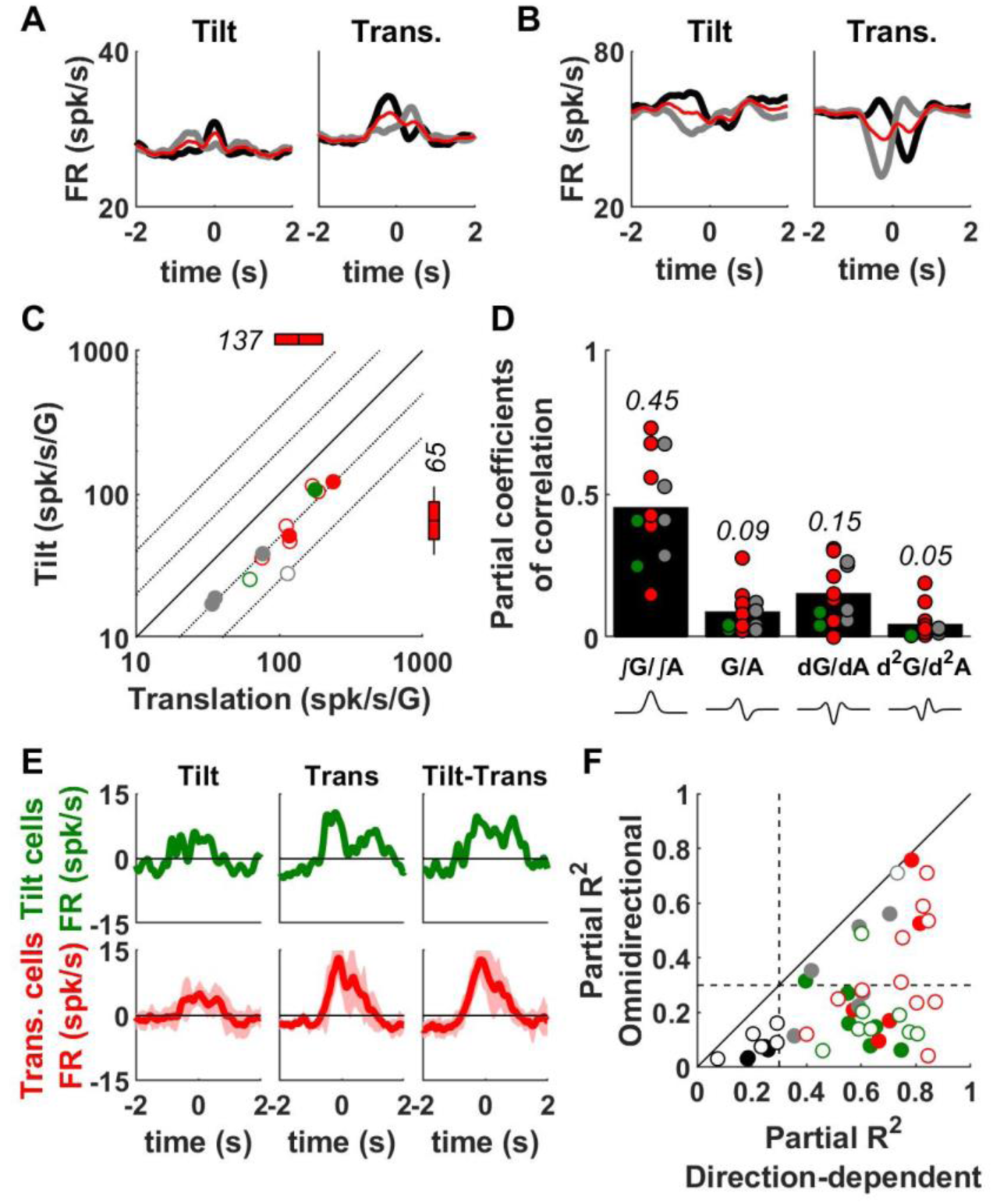
Omnidirectional modulation. In general, neuronal responses reverse when stimulus direction is reversed, but this is not typically the case with vestibular neurons tuned to translation, which have a large contribution of omnidirectional tuning (Laurens et al., 2017). The direction-dependent neuronal modulation can be visualized by computing the difference in firing rate between opposite motion directions (ΔFR, see Methods). In contrast, averaging the firing rate across motion directions reveals an omnidirectional response (see Methods). Omnidirectional responses are evaluated using the same statistical approach as direction-dependent responses (see Methods). A total of 13/46 cells exhibit significant omnidirectional responses (**Suppl. Table 2**). Their properties are summarized here. **(A)** Firing rate of a translation-selective cell during tilt and translation, along the PD (black) and anti-PD (grey). The average of these two curves (red) exhibits a positive omnidirectional response during translation. **(B)** Firing rate of another translation-selective cell exhibiting a negative omnidirectional response. **(C)** Peak-to-trough amplitude of the omnidirectional responses during tilt versus translation. The color code indicates the cells’ classification based on direction-dependent responses (green: tilt-selective, red: translation-selective, gray: composite). Positive and negative omnidirectional modulations are indicated by filled and open symbols, respectively. Omnidirectional modulation is larger during translation than during tilt (p = 2.10^-4^, paired Wilcoxon test across all cells, n=13, with significant omnidirectional responses). The boxes and whiskers represent the geometric mean, confidence interval (boxes) and standard deviation (whiskers) of the response gain of translation-selective cells (other cells types are not shown due to the low number of responsive cells). **(D)** The omnidirectional response component followed mostly ∫G/∫A dynamics. At the population level (tilt-, translation-selective and composite cells pooled), the partial coefficient of correlation of the ∫G/∫A component was higher than that of other components (multiple Wilcoxon signed rank test, Bonferroni correction, p <10^-3^, n = 13). **(E)** Average omnidirectional response in tilt-(n=2) and translation-selective (n=7) cells. The sign of the modulation was inverted prior to averaging in cells where the modulation is negative. In agreement with panels (C) and (D), the modulation is higher during translation than tilt and follows a monophasic profile characteristic of the ∫G/∫A dynamic component (**Fig. S1**). Note that the modulation does not reverse during tilt-translation compared to translation, even though the direction of the translational stimulus is reversed during tilt-translation compared to translation. This is expected since omnidirectional modulation is not affected by stimulus direction. **(F)** The partial R^2^ of the direction-dependent firing rate modulation was higher than that of omnidirectional modulation in all cells. Data from all recorded cells (n=46) are shown. The color code indicates the cells’ classification based on direction-dependent responses (see **Suppl. Table 2**; green: tilt-selective, red: translation-selective, gray: composite, black: NR). The broken black lines indicate the threshold of 0.3, below which direction-dependent or omnidirectional responses are not considered significant.

**Figure 4 Supplement 1:**
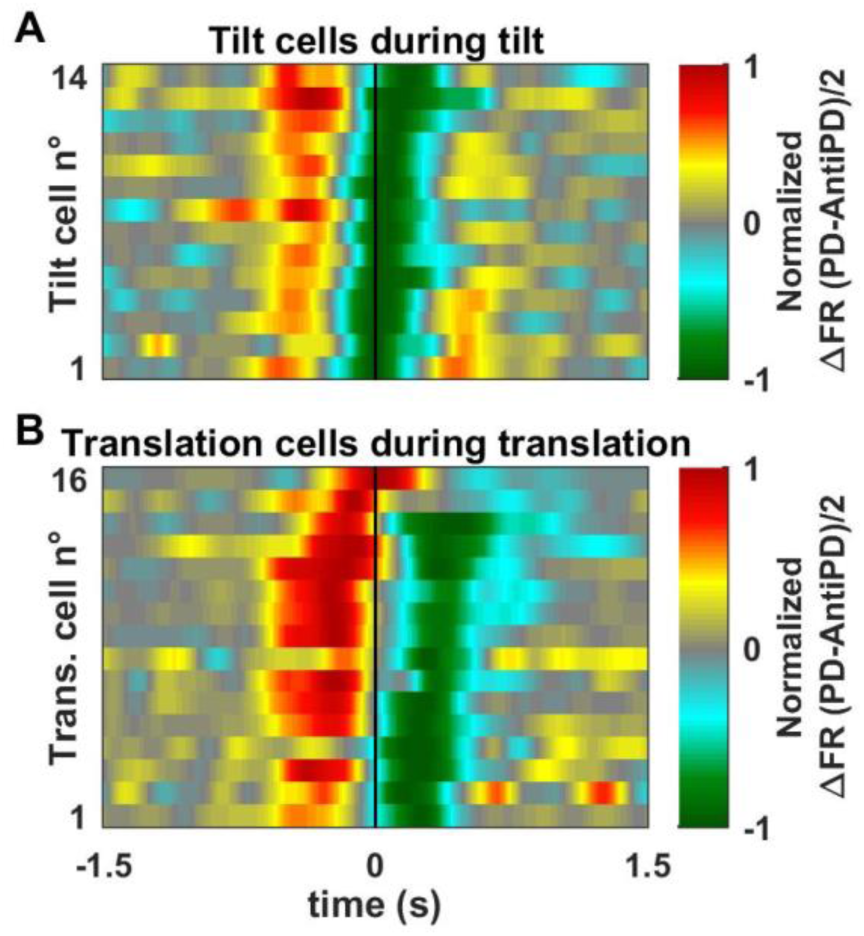
Population response of (A) tilt- and (B) translation-selective Purkinje cells. We computed the direction-dependent firing rate modulation (ΔFR) of tilt-selective cells (during tilt) and translation-selective cells (during translation). The modulation was normalized with respect to its peak absolute value. Neurons were ordered according to their response timing (timing of the negative peak in tilt-selective cells, average between the timing of the positive and negative peaks in translation-selective cells). The resulting population responses are represented using an intensity scale.

**Figure 4 Supplement 2:**
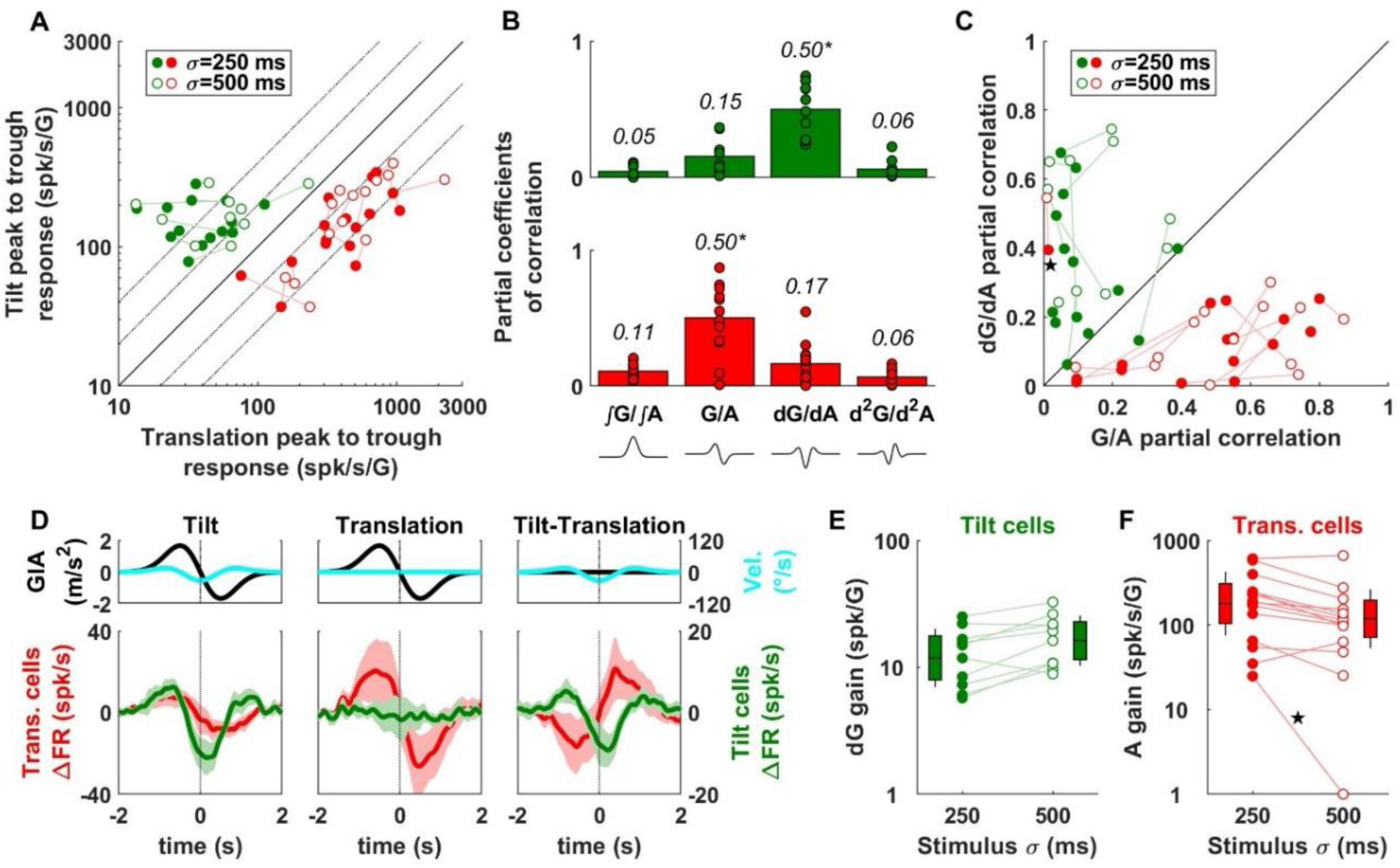
Comparison of neuronal responses during transient motion of longer duration. We recorded the responses of n=10 tilt-selective and n=14 translation-selective cells during transient motion with σ=500ms. **(A)** Peak-to-trough modulation during tilt versus translation with σ=250ms and σ=500ms (open and filled symbols respectively). Data points from individual neurons are joined by lines. **(B)** Partial coefficients of correlation of the dynamic components in response to the σ=500ms stimuli. The dG/dA component has a higher partial coefficient of correlation compared to all other components (p<0.001, multiple paired Wilcoxon tests, Bonferroni correction) in tilt cells. The G/A component has a higher coefficient in translation cells (p<=0.02). These results are identical to those observed with the σ=250ms stimuli (Fig. 3B). **(C)** Cell-by-cell comparison of the partial correlation of the G/A and dG/dA dynamic components (same symbols as in A). **(D)** Average responses ΔFR of tilt (green) and translation (red) cells in response to the σ=500ms stimuli. **(E)** dG response gains of tilt cells, shown for both σ=250ms and σ=500ms stimuli. Boxes and whiskers indicate the geometrical mean, CI and SD (with σ = 250 ms: mean = 12 spk/G, CI=[8 - 18]; with σ = 500 ms: mean = 16 spk/G, CI=[11 - 23]). Although the confidence intervals overlap, there was a small gain increase in all cells resulting in a significant increase at the population level (paired Wilcoxon test, p = 0.02). **(F)** A (acceleration) gain of translation-selective cells. The cell marked by a star exhibits an atypical response pattern and is excluded from the statistical analysis. Boxes and whiskers indicate the geometric mean, CI and SD (with σ = 250 ms: mean = 180 spk/s/G, CI=[104 - 308]; with σ = 500 ms: mean = 118 spk/s/G, CI=[71 - 195], p = 0.03, paired Wilcoxon test).

**Figure 5 Suppl. 1:**
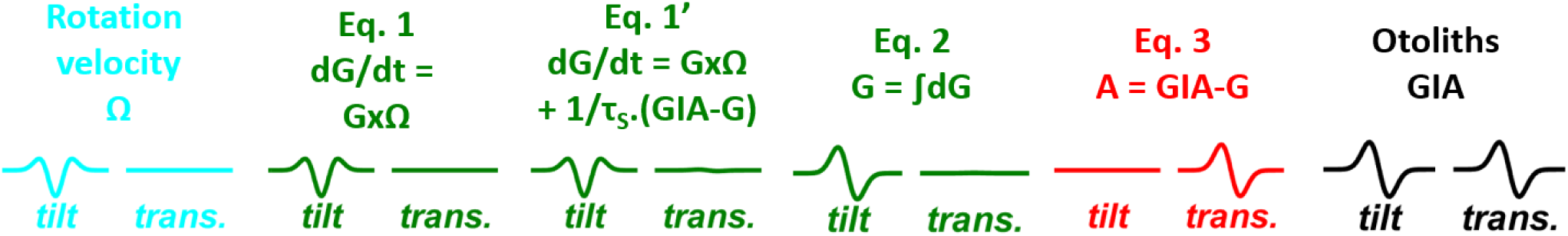
Summary of the motion variables and equations used in **Fig. 5**, and illustration of the temporal profiles of these variables during tilt and translation (σ=250ms). Note that the somatogravic feedback (converting ***eq. 1*** into ***eq. 1’***) has not been considered in the analysis of neuronal data recorded here as it plays a minor role in the frequencies used. Specifically, during passive motion, an otolith feedback signal (equal to (GIA-G)/τ_S_, see Laurens and Angelaki 2011, 2017) is added to ***eq. 1*** to compute tilt velocity, resulting in eq. 1’: dG/dt = GxΩ + (GIA-G)/τ_S_, where τ_S_ is a time constant of ̃0.5s.

**Supplementary Table 1:** Classification of cells into tilt-, translation-, GIA-selective and composite. Neurons were classified independently based on sinusoidal motion (0.5Hz, ±0.2G, Angelaki et al. 2004, Laurens and Angelaki 2013) and transient motion. The results of both classifications are presented as a contingency table. The classification obtained using both data sets was identical for 28/46 (61%) cells. Out of the remaining 18 cells, 14 were classified as tilt-, translation- or GIA-selective based on sinusoids and composite (n=6) or non-responsive (n=8) cells based on transients. Cells are classified as composite when none of the tilt, translation and GIA model is significantly higher than the others and as non-responsive when the signal to noise ratio (measured as the VAF of the composite model) is low. Therefore, the change in the classification of these cells may be explained by the lower amplitude (0.1G) of the transient motion that weakens neuronal responses as well as the statistical power of this stimulus. The present study used the more conservative classification based on transients. We verified that changing the classification scheme did not alter the main conclusions. Purkinje cells classified as GIA-selective or composite (Laurens et al 2013b) have not been further considered here.

|  |  | Classification based on transients |  |  |  |  | Total |
| --- | --- | --- | --- | --- | --- | --- | --- |
|  |  | Tilt | Trans. | GIA | Comp. | NR |  |
| Classification based on sinusoids | Tilt | 12 | 0 | 0 | 1 | 5 | 18 |
|  | Trans. | 0 | 14 | 0 | 5 | 1 | 20 |
|  | GIA | 2 | 1 | 0 | 0 | 2 | 5 |
|  | Comp. | 0 | 1 | 0 | 1 | 0 | 2 |
|  | NR | 0 | 0 | 0 | 0 | 1 | 1 |
|  | Total | 14 | 16 | 0 | 7 | 9 | 46 |
| *classification used in the manuscript* |  |  |  |  |  |  |  |

**Supplementary Table 2:** Classification of cells based on omnidirectional responses compared to direction-dependent responses. The results of both classifications are presented as a contingency table. Few (13/46, 28%) cells, mainly translation-selective, exhibit significant omnidirectional responses. Most (11/13, 85%) omnidirectional responses occur specifically during translation.

|  |  | Classification based on direction-dependent responses |  |  |  |  | Total |
| --- | --- | --- | --- | --- | --- | --- | --- |
|  |  | Tilt | Trans. | GIA | Comp. | NR |  |
| Classification based on omnidirectional responses | Tilt | 0 | 0 | 0 | 0 | 0 | 0 |
|  | Trans. | 1 | 7 | 0 | 3 | 0 | 11 |
|  | GIA | 0 | 0 | 0 | 0 | 0 | 0 |
|  | Comp. | 1 | 0 | 0 | 1 | 0 | 2 |
|  | NR | 12 | 9 | 0 | 3 | 9 | 33 |
|  | Total | 14 | 16 | 0 | 7 | 9 | 46 |
|  |  | classification used in the manuscript |  |  |  |  |  |

